# Neuroepithelial organoid patterning is mediated by Wnt-driven Turing mechanism

**DOI:** 10.1101/2021.01.11.426254

**Authors:** Abdel Rahman Abdel Fattah, Sergei Grebenyuk, Idris Salmon, Adrian Ranga

## Abstract

Cell patterning in epithelia is critical for the establishment of tissue function during development. The organization of patterns in these tissues is mediated by the interpretation of signals operating across multiple length scales. How epithelial tissues coordinate changes in cell identity across these length scales to orchestrate cellular rearrangements and fate specification remains poorly understood. Here, we use human neural tube organoids as model systems to interrogate epithelial patterning principles that guide domain specification. *In silico* modeling of the patterning process by cellular automata, validated by *in vitro* experiments, reveal that the initial positions of floor plate cells, coupled with activator-inhibitor signaling interactions, deterministically dictate the patterning outcome according to a discretized Turing reaction-diffusion mechanism. This model predicts an enhancement of organoid patterning by modulating inhibitor levels. Receptor-ligand interaction analysis of scRNAseq data from multiple organoid domains reveals WNT-pathway ligands as the specific inhibitory agents, thereby allowing for the experimental validation of model predictions. These results demonstrate that neuroepithelia employ reaction-diffusion-based mechanisms during early embryonic human development to organize cellular identities and morphogen sources to achieve patterning. The wider implementation of such *in vitro* organoid models in combination with *in-silico* agent-based modeling coupled to receptor-ligand analysis of scRNAseq data opens avenues for a broader understanding of dynamic tissue patterning processes.

## Introduction

During embryonic development, complex tissue morphologies emerge from rearrangements of simpler modules. One such module is the single-cell-thick epithelial sheet, which, through precise and timely patterning and morphogenetic events, allows for the specification of cellular identity and tissue growth. To orchestrate the morphology of epithelia in time and space, tissue-wide spatial reference frames are established by biochemical and mechanical signals^1^, which are interpreted by individual cells to determine fate decisions and ultimately result in short and long range intercellular interactions. The specification of cell fate in the developing neural tube presents a canonical illustration of epithelial pattern formation, where multiple domains are dynamically specified in response the strength, exposure time and cross talk between diffusible signaling factors such as sonic hedgehog (SHH), bone morphogenic protein 4 (BMP4) and retinoic acid (RA). Our understanding of these patterning processes have thus far largely relied on animal models which, while highly reproducible and predictive, nonetheless comprise multiple layers of feedback and redundancy, making it challenging to uncover the underlying regulatory mechanisms.

Due to their capacity for *in vivo*-like multicellular organization^2^, organoids have emerged as tractable *in vitro* platforms to study patterning and morphogenesis in tissues as varied as the optic cup^3,4^, the intestine^5,6^, and the cortical plate^7^, and in gastrulation of the mammalian embryo^8^. Neural tube organoids (NTOs) derived from pluripotent stem cells (PSCs) are a promising model system to explore epithelial patterning, as they have been shown to recapitulate aspects of dorsoventral (DV)^9^ and anteroposterior (AP)^10^ patterning of the neural tube though morphogen stimulation. Given specific morphogen stimulation, NTOs exhibit a floor plate (FP) domain^11^ corresponding to the organizing region of the ventral neural tube and a source of a SHH gradient which encodes DV domain specification. The establishment of this FP domain is not fully explainable by cytoskeleton-mediated symmetry breaking events^11,12^ or by the applied signaling factors, as these cultures lack a notochord, which *in vivo* establishes a reference frame for the initiation of SHH-mediated patterning^9,11,13,14^. This raises the question as to the nature of FP patterning *in vitro*, and how this may relate more generally to the establishment of epithelial patterning.

To describe patterning dynamics, positional information (PI)^15^ as well as reaction-diffusion (RD)^16^ models have been extensively studied *ex vivo*, *in vitro* and *in silico*. While these two principles have been portrayed as contradictory and mutually exclusive^14,17^, recent studies have seen the application of combined PI-RD approaches to explain digit patterning^18^ as well as epithelial NT domain orchestration^19^. These studies suggests that morphogen source position, in combination with the characteristics of interacting diffusible species, are needed to explain epithelial patterning.

Here, we use a human NTO (hNTO) *in vitro* platform and a cellular automaton (CA) *in silico* model to investigate FP patterning dynamics. We establish the model’s fidelity by benchmarking to in vitro experiments and systematically explore the parameter space which is permissive for recapitulating hNTO pattern types and frequencies across various length scales. We also investigate model parameters to predict *in vitro* routes to enhance patterning in hTNOs, and identify concentration of inhibitor species as a powerful modulator of patterning phenotype. An analysis of receptor-ligand interactions of single cell RNA-sequencing (scRNA-seq) data is used to identify the specific inhibitor species. Finally, we perform perturbation experiments to validate model predictions. The use of simple and tunable CA model in combination with analysis of transcriptomic data reveals insights into mechanisms of human neural tube patterning, and may be extendable to other epithelia.

### Human neural tube organoids: an *in vitro* platform for epithelial patterning

In order to study patterning processes in epithelial tissues, we rely on a highly defined *in vitro* platform in which single hPSCs are differentiated into pseudostratified epithelial hNTOs within a synthetic polyethylene glycol (PEG) hydrogel matrix supplemented with laminin^20^ (**Fig 1a**). In the absence of signaling factors, these organoids remain largely unpatterned and have a forebrain identity. To initiate a FP domain, organoids are treated with RA to caudalize the default forebrain identity^11,12^ and with smoothened agonist (SAG) to provide a ventralizing signal through the activation of the SHH pathway^9^. By day 11 of differentiation, hNTOs display various FP expression phenotypes, as identified by the FP marker FOXA2 (**Fig 1a**). These could be categorized into three groups: those that exhibited a patterned FP, a scattered FP, and those that were devoid of the fate. FOXA2+ hNTOs represented ~60% of all organoids, of which ~34% were patterned (**Fig 1b**). FP patterning in these organoids was initiated after RA-SAG treatment by the emergence of FOXA2+ cells. These cells were initially scattered within the organoid in a largely disorganized manner, and over time rearranged such that some FOXA2+ domains gradually underwent patterning^21^ (**Supplementary Fig 1**). Despite all starting from single cells, organoids were observed to reach different sizes by day 11. No significant difference in organoid size was observed between patterned and scattered organoids (**Fig 1c**), and average patterning frequency was independent of organoid size (**Fig 1d**), suggesting that the occurrence of patterning in this system is independent of domain size.

**Figure 1.**
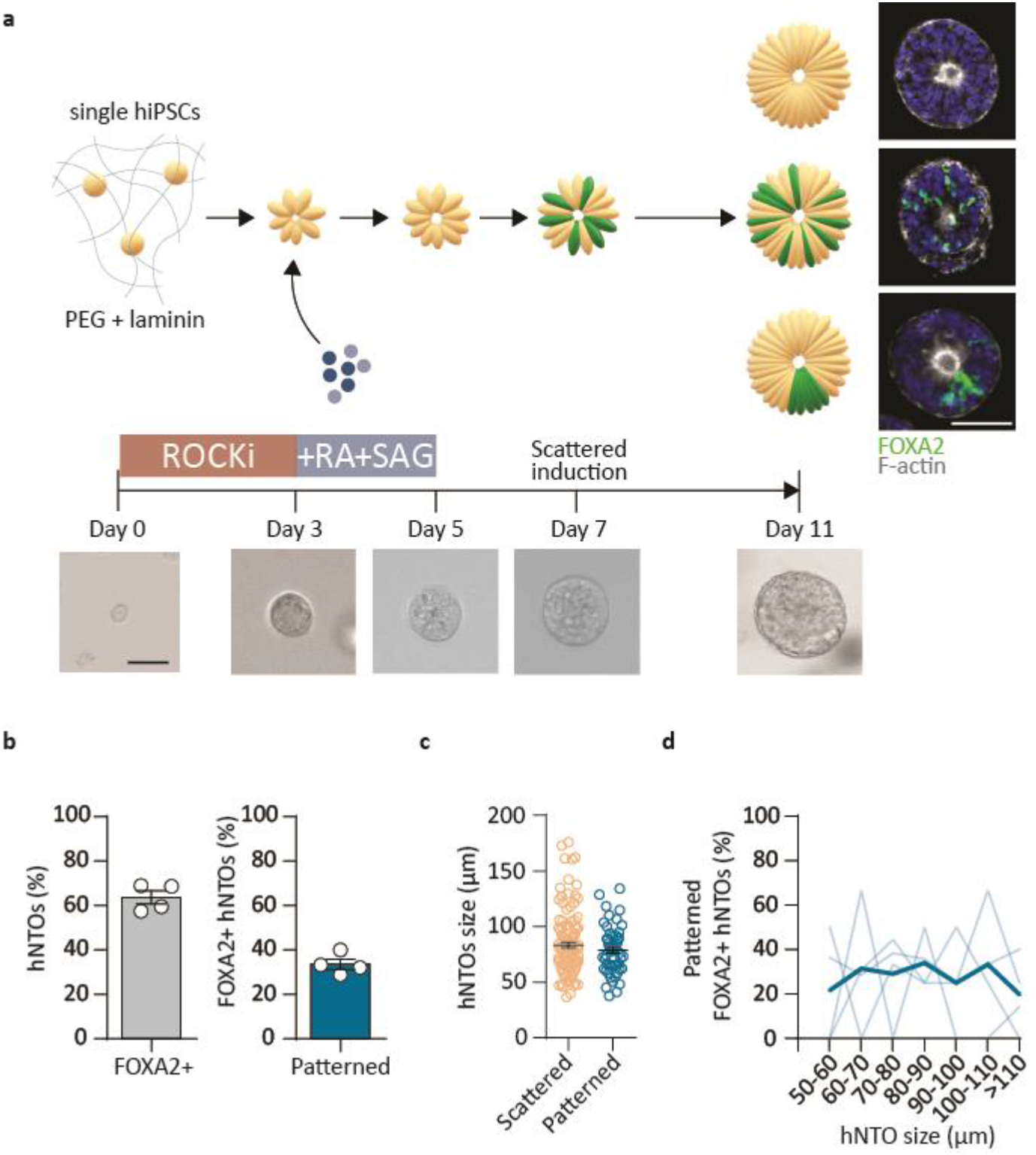
Human neural tube organoid floor plate characterization. **a** Organoids derived from single hiPSCs in synthetic matrices result in patterned, scattered or negative FP expressions. Representative brightfield images of hNTOs at various timepoints, with representative expression of FOXA2 and F-actin at day 11 endpoint (nuclei: Hoechst). **b** FP induction and patterning frequencies (n = 4, 269 hNTOs assessed). **c** Quantification of organoid size (diameter) for scattered and pattered hNTOs. **d** Quantification of patterning efficiency as a function of organoid size (dark blue line represents average at each size range, light blue lines represent individual replicates). Scale bars: 50 μm.

### 1-D Turing model for patterning human neural tube organoids

To explore mechanisms that could explain FP patterning in hNTOs we sought to develop an *in silico* model that could best describe *in vitro* observations. We reasoned that a RD approach^16^ could be suited to model the transition between an initially scattered expression profile towards a gradually patterned domain based on previous approaches which have used RD to describe patterning in peri-gastrulation tissues^22^ and retinal organoids^23^ *in vitro*.

To tailor the model to our hNTO, we relied on a simple 1-D activator-inhibitor RD system with second-order production rates and first-order decay terms, similar in general form to those previously used in other organoid model systems^16,24–26^ (**Supplementary Fig 2a**). Periodic boundary conditions were implemented in order to mimic the epithelial morphology of hNTOs^9,20^, and parameters of the partial differential equations were chosen based on those previously applied for RD organoid models to describe the production and decay rates of morphogens^23^ (**Supplementary Fig 2a**). Cells in the domain could be either FOXA2+, represented as source cells (SCs), or FOXA2-, represented as inactive cells (ICs). To simulate the initially scattered FP expression, a random perturbation was imposed on the activator signal (Φ_a,m_). Once perturbed, the reaction between activator and inhibitor species led to the assignment of each cell in the model as a SC wherever the associated Φ_a,m_ was larger than the average overall activator signal Φ_a,avg_, or as an IC where Φ_a,m_ was lower than Φ_a,avg_ (**Supplementary Fig 2b**). The calculated expression profiles across the domain at each time step are deterministic in this model, with this RD system converging into a specific pattern for a given set of parameters such as domain size, and activator and inhibitor diffusion coefficients D_a_ and D_i_.

It is known that domain size plays an important role in RD systems, with patterning typologies being domain size dependent^24,25^. Indeed we observed that an increase in domain size resulted in larger number of SC poles (**Supplementary Fig 2c**), indicating that the model led to pattern size-dependence which contradicted *in vitro* observations. Most importantly, we observed significant phenotype heterogeneity *in vitro*, with a mix of patterned, scattered and unpatterned organoids within the same culture, whereas this *in-silico* model rather functioned as an “all-or-nothing” switch, resulting in either fully patterned or fully negative (i.e. unpatterned with no SCs remaining) phenotypes for a given set of physical parameters (**Supplementary Fig 2c**). Altogether, this suggests that a canonical RD approach cannot fully recapitulate observed *in vitro* expressions. This led us to investigate modeling approaches that could allow for SC expression profiles similar to those seen in-vitro, including varied patterning phenotypes, heterogeneous phenotypes within an organoid population, and where these findings would be domain-size independent.

### Cellular automata model recapitulates *in vitro* floorplate expression heterogeneity

In our model the only source of stochasticity that can lead to expression heterogeneity is the initial perturbation and location of SCs. We therefore considered the use of a cellular automaton^27^ (CA) model, which would allow to specify the initial positions of SCs which drive the evolution and dynamics of the system. We implemented an *in silico* 1-D elementary CA model (**Fig 2a**), where simulations are run for ten time steps and the identity of each unit cell (SC or IC) in the domain is updated at each time step. Here, SCs are allowed to emit fate-activating and fate-inhibiting signals with concentrations Φ_a_ and Φ_i_ respectively (**Fig 2a**). The change in the concentration of these signals after every time step is governed by the equation Φ = Be^−λx^, where B represents a constant morphogen concentration maintained through replenishment at the source, and decay constant λ represents morphogen diffusion characteristics through the tissue. Together, B and λ govern the interaction range of SCs with their cellular neighborhoods (**Fig 2a**). Cells in which the net resulting interaction between activator and inhibitor (Σ(Φ_a,n_ - Φ_i,n_)) is less than a minimum threshold become (or remain) ICs, whereas those above the threshold become (or remain) SCs. Survivor cells are by default SCs and, when surviving through all ten steps, contribute to the final SC expression, which can then be classified as patterned, scattered or negative (**Fig 2a**).

**Figure 2.**
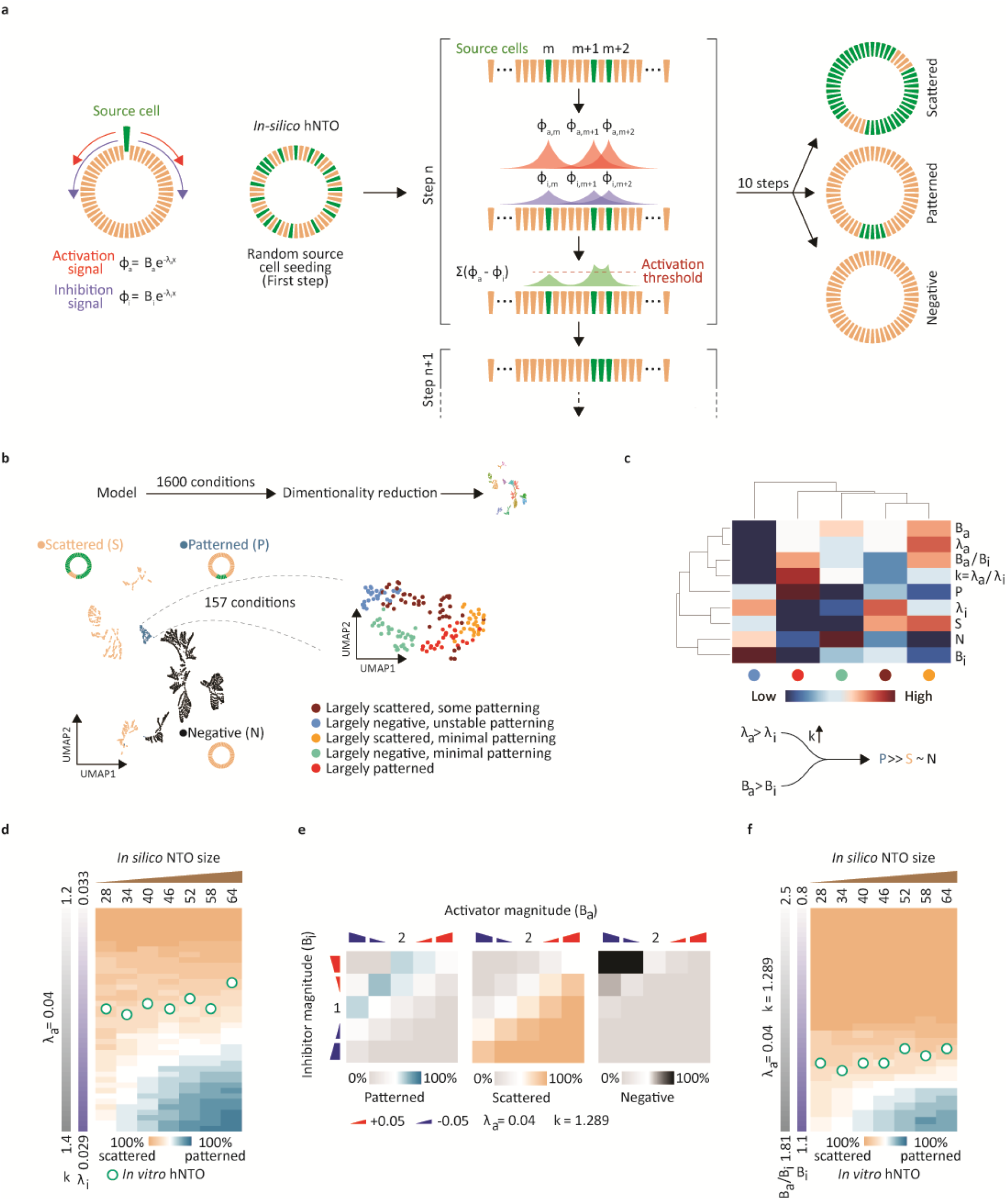
*In silico* modeling of FP patterning in NTOs. **a** A cellular automata model combines morphogen source position and associated diffusion characteristics. SCs emit activation and inhibition signals and interact with adjacent cells over 10 steps resulting in various SC expressions. **b** Combinatorial parametric analysis of *in silico* 1600 conditions, with 157 conditions that result in patterning expressions of SCs. **c** Hierarchical clustering of parametric analysis output for patterning conditions. **d** Mapping *In vitro* pattering efficiencies (white and green circles from **Fig 1d**) across various sizes to *in silico* data for various values of k and λ_a_ = 0.04. **e** SC expression output as a function of various activator and inhibitor magnitudes. **f** Mapping *in vitro* pattering efficiencies (white and green circles from **Fig 1d**) across various sizes to *in silico* data for various values of inhibitor magnitudes, k = 1.289 and λ_a_ = 0.04. (for every simulated data point, n = 3 for a total of 1,500 *in silico* NTOs)

We hypothesized that the *in vitro* expression of individual cells and the ensuing generation of domains could be minimally described in terms of concentration (B_a_, B_b_), as well as the diffusion characteristics (λ_a_, λ_b_) of activation and inhibition signaling molecules. We therefore combinatorially explored parameters of B and λ for both inhibitor and activator to identify conditions that could lead to patterning of SCs. Out of 1,600 conditions analyzed, only 157 led to SC patterning (**Fig 2b**). Notably, the highest patterning frequencies (red cluster in **Fig 2b**) occurred when the activator diffusivity was lower than that of the inhibitor species (in terms of decay constants: λ_a_ > λ_i_) (**Fig 2c**).

To test the CA model in a neural tube-specific context, we chose λ_a_ = 0.04, reflecting the observed decay length values for SHH *ex vivo*^28^, and found that patterning occurred for k > 1, where k = λ_a_/λ_i_ is the decay constant ratio (**Supplementary Fig 3**). These are conditions within the short range activator and long range inhibitor regime, representing Turing instability scenarios known to favor patterning during morphogenesis^16^. Therefore the CA approach we develop here can be interpreted as a discretized Turing patterning model, which has as a defining feature the ability to capture specific SC expression patterns given a fixed set of parameters (**Supplementary Fig 4**).

By mapping the *in vitro* rate of patterned organoids to model outputs we found the specific ratio between activator and inhibitor decay constants (k ≈ 1.29) which best describes the heterogeneity observed *in vitro* (patterning events ~34%) (**Fig 2d**). This indicates that our CA model captures both specific organoid phenotypes, as well as their heterogeneous distribution within a population, suggesting that FP patterning in these epithelial organoids follows a discretized Turing patterning approach, as predicted by our CA model.

As our model relies on previously reported morphogen diffusion values derived from *in vivo* experiments, we sought to compare the diffusion dynamics of hNTOs to previously reported *in vivo* observations. We therefore performed fluorescence recovery after photobleaching (FRAP) experiments on day 5 hNTOs, which best describe the microenvironmental state at the onset of patterning (**Supplementary Fig 5**) and found a diffusion coefficient of D_FL_ = 61.8 μm^2^.s^−1^. This result confirmed that hNTOs have similar diffusion dynamics as those reported *in vivo* (D_FL_ ~ 50 - 100 μm^2^.s^−1^) ^29 30^.

An important characteristic of this model is its ability to demonstrate size-independent patterning, where the frequency of pattering remains relatively constant across different domain sizes (**Supplementary Fig 3**). This allowed the CA model to predict *in vitro* FP expressions across various length scales where the average change in patterning frequency given a change in organoid size was negligible (**Supplementary Fig 6a**). Notably, the model matched FP domain size trends in scattered and patterned hNTOs (**Supplementary Fig 6b)**, suggesting that this modeling approach recapitulates the dynamics leading to each expression type. Moreover, the CA model can not only discriminate between scattered and patterned expressions, but also between patterning subtypes. Here, we accurately recapitulated the relatively rare 2-pole and 3-pole pattern subtypes (**Supplementary Fig 7a**), with *in vitro* observations of pole location and frequency quantitatively matching *in* silico predictions (**Supplementary Fig 7b, c**).

This model predicts that patterning frequencies can not only be changed by varying molecule diffusion characteristics λ, which are difficult to modulate *in vitro*, but equally through activator or inhibitor magnitudes (**Fig 2e**), which can be readily manipulated through pharmacological perturbations. Indeed, by fixing the diffusion characteristics of activator and inhibitor molecules, the model predicted that an increase in patterning frequency is possible through an increase of the inhibitor magnitude Bi (**Fig 2f**).

### Receptor-ligand interaction analysis reveals WNT as putative inhibitor of the RD system

In order to identify the inhibitor species in the hNTO system, we performed receptor-ligand interaction analysis^31^ on an scRNAseq dataset obtained from dissociated hNTOs at day 5, corresponding to hNTOs after RA-SAG exposure but before FP induction, and at day 11, after FP induction and patterning^20^. From this dataset, only those cells which could be identified as having dorsal (D), intermediate (I) or ventral (V) fates were retained, and were represented on UMAPs highlighting their states at different time points (**Fig 3a**) and by D-I-V assignments (**Fig 3b**). These cells also displayed domain-specific hallmark genes corresponding to expected *in vivo* expression profiles (**Fig 3c**). We next focused our analysis on understanding the interactions between cells in the D-V domains, which represent the known morphogen sources in the neural tube, and between cells in the V-V domain, which represent domain self-interaction with the potential to highlight inhibitory molecule candidates.

**Figure 3.**
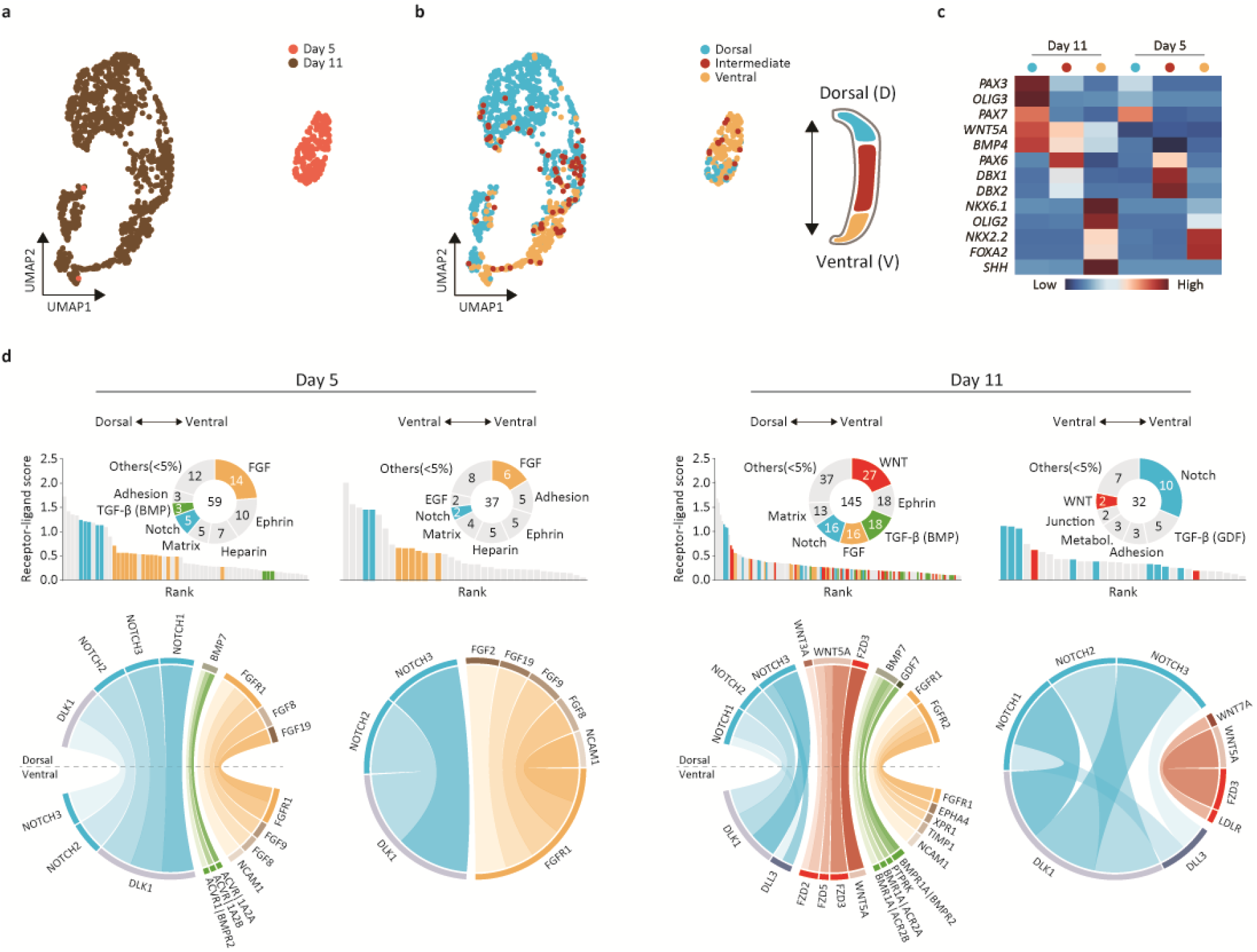
Receptor-ligand interaction in hNTOs. **a** UMAP highlighting day 5 and 11 of D, I, and V hNTO cells color coded by days. **b** UMAP color coded by domain identity D, I, and V. **c** Selected marker genes for dorsoventral identities. **d** Receptor-ligand analysis of day 5 and day 11 hNTOs. Interactions belonging to NOTCH, FGF, BMP and WNT signaling pathways are color coded blue, yellow, green, and red respectively.

This analysis revealed 59 and 37 significant interactions for D-V and V-V respectively on day 5, compared to 145 and 32 interactions for D-V and V-V respectively on day 11 (**Fig 3d**). We further grouped these interactions and categorized them by their major participating ligand or receptor, and highlighted only these interactions that involve known modulators of the FP and ventral domains including antagonists (WNT^32,33^ and BMP^34^) and agonists (NOTCH^35,36^ and FGF^37,38^) (**Fig 3d**).

At day 5, we observed FGF, NOTCH, and BMP activity in D-V interactions and FGF and NOTCH in V-V ones (**Fig 3d**). FGF activity was transient as its fraction declined from day 5 to 11, which is in line with a previous *in vivo* report of transient FGF signaling being required for FP induction^34^. The presence of BMP suggested that dorsal cells, although representing only a small population at day 5, already played at this early time point a critical role as sources of BMP required for DV patterning. The top 5 NOTCH activities at day 5 all involved *DLK1*, a known inhibitor of the NOTCH pathway^35^. After the induction and patterning of FOXA2 at day 11, *DLL3*, another inhibitor of NOTCH, was also involved in both interacting pairs. The introduction of WNT signaling in both D-V and in V-V interactions at day 11 represents a significant change in the receptor-ligand makeup compared to day 5. This suggests that WNT signaling and not NOTCH, FGF, or BMP is a marker of a maturing organoid characterized by FP induction and patterning. Furthermore, since FOXA2+ hNTOs are largely devoid of dorsal cells^20^, it is likely that WNT, and not BMP from dorsal cells, represents the inhibition signaling within single organoids.

### WNT signaling promotes floorplate patterning through inhibition

Our CA model predicted that changes to the abundance of inhibitory molecules would affect patterning frequency (**Fig 2f**). In order to test this prediction experimentally with our newly identified putative inhibitor, we chose to modulate WNT signaling in our cultures using CHIR99021, a small molecule WNT activator (via GSK3β inhibition). We supplemented the growth medium with GSK3βi at different concentrations ranging from 0 to 4 μM (**Fig 4a**) from day 7, immediately after the induction of the FP, until endpoint day 11 (**Fig 4a**). Increasing WNT activity resulted in a decrease in the frequency of the scattering phenotype, an increase in the negative (i.e. no FOXA2+ cells) phenotype, and a rise (up to 2uM) then decline in the patterned phenotype (**Fig 4a**). These *in vitro* values matched with our CA model, with the increase in GSK3βi concentration from 0 to 4 μM corresponding to an increase in inhibitor magnitude Bi from 1 to 1.14.

**Figure 4.**
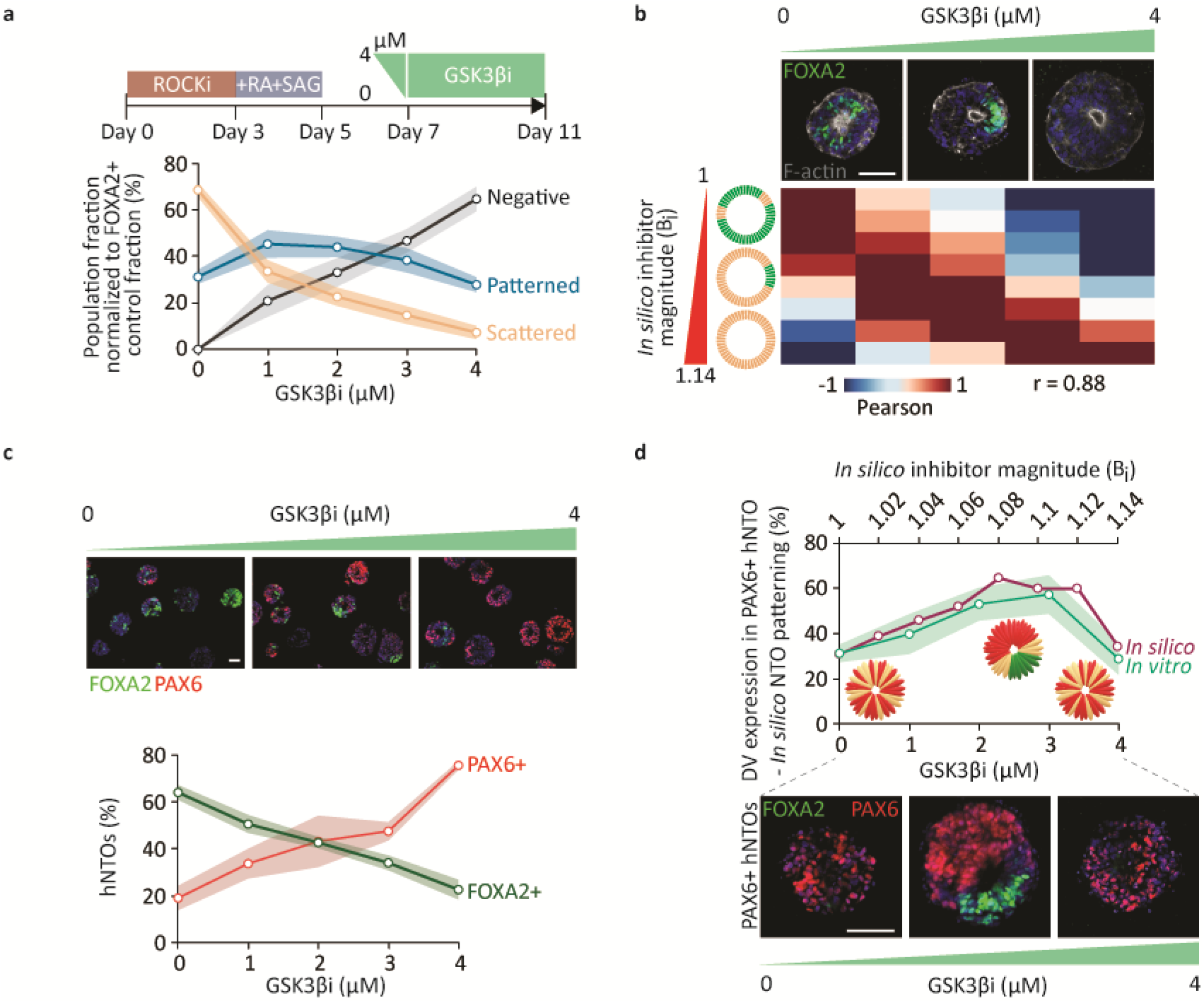
WNT activation affects FP patterning. **a** Scattered, patterned and negative event frequencies for various concentrations of GSK3βi (n = 4 for a total of >300 hNTOs). **b** Correlation analysis between *in vitro* observations for varying concentrations of GSK3βi and *in silico* results for varying inhibitor magnitudes (Pearson r = 0.88), (n = 3 for a total of 1,500 *in silico* NTOs per data point and *in vitro* data from **a**). **c** Effect of WNT activity on FOXA2 and PAX6 expressions (n = 4 for a total of >300 hNTOs). **d** Trend comparison of DV patterning in FOXA2+/PAX6+ hNTOs for various concentrations of GSK3βi and *in silico* patterning for various inhibitor magnitudes (*In silico* data from **b** and *in vitro* data from **c**). Scale bars: 50 μm.

We hypothesized that such an increase in patterning events would be accompanied by a larger domain diversity, including the presence of more dorsal domains within the same organoid, which would better recapitulate *in vivo* conditions. We therefore interrogated the presence of more dorsal identities such as PAX6, and found that increasing GSK3βi concentrations resulted in a gradual decrease of FOXA2+ hNTOs, with a converse increase in PAX6+ organoids (**Fig 4b**). This is in line with the reported promotion of *in vitro* dorsal identity in the neural tube by WNT activation^34^. Furthermore, with increasing GSK3βi concentrations, we found that the fraction of dorsoventral patterning in PAX6+ hNTOs follows a similar trend to *in silico* patterning (**Fig 4c**). This underscores the close link between FP patterning and dorsoventral patterning events, and, in the context of organoids, argues for a control mechanism where multiple domains coexist within the same organoid when secondary sources of inhibition or activation are created as a result of domain-domain interactions^28,35^.

Due to the increase in dorsal fates upon exposure to GSK3βi, and since dorsal fates can be sources of WNT secretion (**Fig 4d**), we next thought to investigate the effect on expression trends with ICs as surrogates for dorsal cells and as secondary sources of inhibition signals (**Supplementary Fig 8a**). When comparing the contributions to inhibition signaling from SC (x1) and IC (x2), we found that the best correlation to *in vitro* data were achieved in conditions where SCs were the sole sources of inhibition (**Supplementary Fig 8b,c,d**). This *in silico* result suggests that FP patterning does not require dorsal domains. At first this seemed contradictory to our previous observation (**Fig 4c**) given the close relation between FP patterning and dorsoventral patterning events with increasing WNT activity. However, we reason that upon WNT activation, FP patterning events are increased through enhanced inhibitor interactions, but that concomitantly, WNT exposure promotes PAX6 expression in FOXA2-regions in the organoid, resulting in dorsoventrally patterned hTNOs. Therefore, our model setup predicted that while FP patterning did not require dorsal domains, dorsal domain patterning depended on patterned FP expressions. Indeed, hNTOs with patterned FPs at a GSK3βi concentration of 0 μM were largely (~66%) devoid of the PAX6 fate (data not shown), further underscoring that FP patterning does not require, neither does it necessarily result in, PAX6 expression within the same organoid. The results, therefore, argue that FP patterning is a self-regulated process as predicted by the model.

Finally, after exploring the upregulation of WNT activity, we tested whether the converse downregulation of WNT activity would have an opposite effect on patterning. The IWP2 small molecule (PROCN inhibitor) was used to decrease WNT activity, resulting in decreased FP patterning frequencies (**Supplementary Fig 9a**) as predicted by the model (**Fig 2e,f**). A concentration of 2 μM of IWP2 correlated best with B_i_ = 0.96, which corresponds to a reduction in inhibitor magnitude (**Supplementary Fig 9b**).

Altogether *in silico* and *in vitro* results suggest that WNT activity plays an important role in shaping FP expression in hNTOs, in accordance with an underlying spatially discretized RD-based model of the system. Importantly, our model predicted that epithelial domains could be rendered largely patterned through an increase of WNT activity, which we verified experimentally. The model suggests that this process occurs by enhancing inhibition signaling. This renders it more difficult for cells to meet the activation threshold which leads to more concise and separated grouping of cells, namely fate polarization or patterns. Increasing WNT activity also allows FOXA2-cells to become PAX6+ which can result in dorsoventral patterning but only at intermediate WNT activities since high levels can eliminate favorable regions, abrogating the FP fate altogether (**Fig 5**).

**Figure 5.**
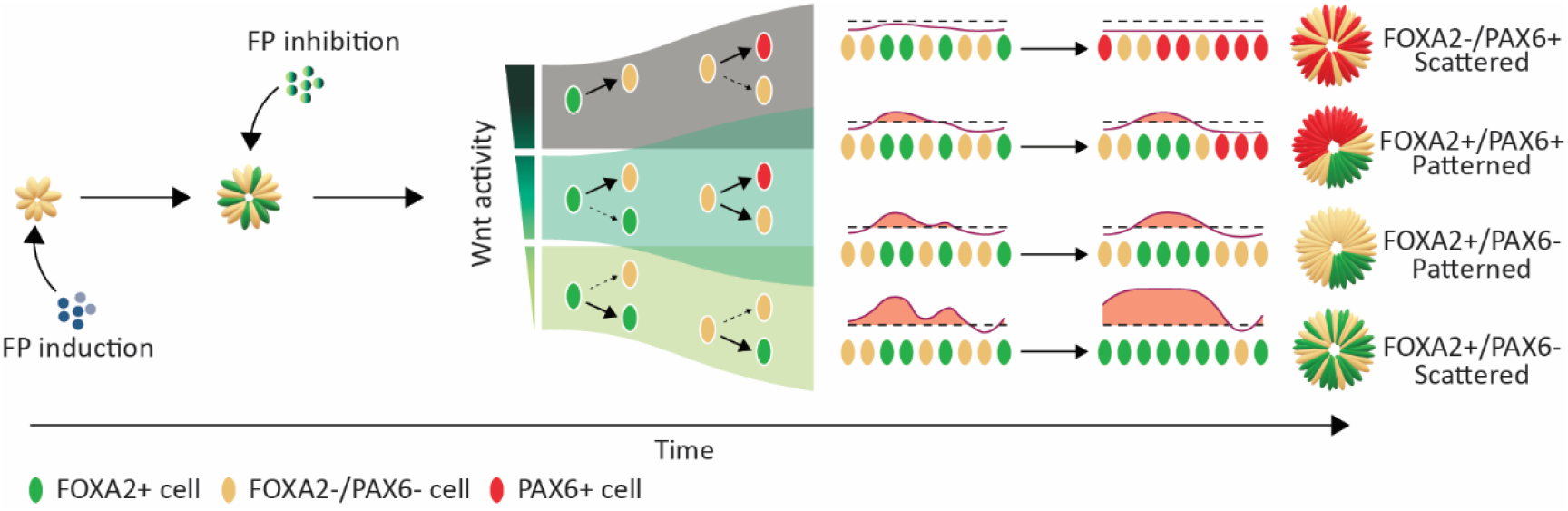
Patterning in epithelial neural tube organoids follow a discretized Turing patterning approach. A model describing epithelial patterning in hNTOs following a cellular automata approach that incorporates morphogen source positions and associated diffusive characteristics. Inhibition modulation though WNT signaling alters final expressions (dashed arrows represent less likely events).

## Discussion

Cell patterning in epithelia is a key event for proper tissue development during early embryogenesis^39,40^. To better understand the underlying mechanisms for the establishment of such patterns, *in silico* models such as PI and RD have been employed^15,16,24^. In a PI approach, cells recognize their position with respect to a source by interpreting the graded morphogens emanating from that source allowing them to pattern according the signal strength and thus their respective distances to the source. This emphasize the importance of morphogen source position in a PI patterning model. By contrast, an RD approach ignores morphogen positions and relies instead on tissue wide secretion and interaction between morphogen species, which allows initially random expressions to converge to predictable pattern configurations. Here, we use an epithelial organoid model of neural tube patterning to show that FP patterning emerges though a discretized Turing mechanism. Specifically, we use a CA-based approach to show that morphogen source position in the tissue and diffusion characteristics of activators and inhibitors control the organization of cells within the domain space, and are thus likely modulators of FP expression in organoids.

Organoids have often been used as simplified models of patterning and have allowed insight into the underlying biochemical^41–44^ and mechanical^1,20,45,46^ factors that control cell fate specification and morphogenesis. We and others have shown that NTOs display patterned FPs^9,11,12,20^ which emerge, remarkably, despite the lack of the notochord, which *in vivo* acts as a spatial reference frame for FP patterning^28,47,48^ through the secretion of SHH morphogens. This suggests that *in vitro* epithelial patterning follows mechanisms that might differ from those employed *in vivo*.

Previous work with epithelial organoids suggests that patterning phenomena can be described solely by an RD approach. For example, as in the case of retinal organoids involving Pax6, Fst, and Tgfb2^23^. This simple gene regulatory network was shown to sufficiently explain how organoids spontaneously self-organize to reflect retinal development. Our study demonstrates that such a canonical form of RD could not explain FP pattering in epithelial hNTOs, as key *in vitro* observations such as patterning heterogeneity and domain size independence could not be recapitulated. Because FP cells are specified by interpreting PI from the notochord SHH source before becoming sources of SHH themselves^28,34^, we hypothesized that a different modelling framework which considered the initial position of FP cells in the RD model would be necessary to better explain our observations.

Models which combine RD with PI have been shown to predict patterning better than either model separately, notably in the context of digit patterning^18^. Here, graded Fgf concentrations (PI) control patterning wavelengths stemming from the interactions (RD) between Sox9, Bmp and Wnt across the domain space, allowing for more robust recapitulation of *in vivo* digit patterning. Furthermore, an *in silico* PI-RD NT model demonstrates how the introduction of intercellular homeoprotein diffusion in the model better orchestrates domain specification by generating more delineated *in vivo*-like domain boundaries^19^.

Our study shows that only when the initial positions of SCs are considered can FP patterns be generated with high fidelity. The developed CA model demonstrated patterning in regimes consistent with a Turing instability mechanism and even exhibited multi-poles patterning as is expected from a RD system. However, only by introducing the activator and inhibitor species at specific SC positions could the reaction between them create highly discretized regions which could persist to maintain a stable pattern. Our results suggest that this discretization adapts the RD system to allow for expression heterogeneity, which ultimately recapitulate *in vitro* FP observations. This model therefore expands on previous models of epithelial patterning and demonstrates that a CA approach can combine the importance of morphogen source position as well as diffusion characteristics to describe epithelial patterning.

Patterning in epithelial tissues rely on complex and dynamic signaling^4,39,49^. In the neural tube^28,34,43,44,48^, WNT^32,33^, BMP^34^, NOTCH^35,36^, FGF^37,38^ and SHH^28^ have been recognized as main modulators of patterning which orchestrate domain specification along the dorsoventral as well as anteroposterior axis. Once these morphogens establish a patterning reference frame, inter-domain signaling assist with domain boundary specification. For example, mediated by SHH, NKX2.2 in the p3 domain prevents the progression of the adjacent pMN domain through OLIG2 inhibition in the ventral NT^28,38^ and in NTOs^9^. Moreover, intra-domain signaling is required for maintenance of cell identity, similar to the role of SHH in the FP^44^. Our modelling results argue that intra-domain signaling is required not only for self-maintenance and activation, but also for self-inhibition. Our receptor-ligand interaction analysis suggests that FP patterning is modulated by WNT, a known inhibitor of ventral identities^33,34,50^. Indeed, FOXA2 abundance variation in response to WNT activity has been previously demonstrated in a microfluidic system^10^, where moderate levels of WNT were necessary for FP induction but higher ones resulted in loss of *FOXA2* and *SHH*. To investigate whether FP patterning is modulated by inhibition, we performed *in silico* modeling and validated the outcome though perturbation experiments to show that increasing FP patterning through inhibitory means is possible, but that high inhibition levels cause loss of FP fate in the majority of cultured hNTOs. We further showed that FP patterning is a self-regulated process that does not require the presence of other inhibitory domains since the best correlation to *in vitro* observations occurs when SCs are the sole sources of inhibition and activation. Interestingly, the expected activator signaling, SHH, was absent in the receptor ligand interaction analysis, which may hint to a different activator *in vitro*. Therefore, while FP induction occurs through a RA-SAG pulse, later pattern maintenance may rely on pathways other than SHH, such as NOTCH signaling, which has recently been shown to regulate ventral domains *in vivo*^35,36^.

Our *in silico* analysis relies on a simplified 1-D CA model without considering cytoskeleton rearrangement^11^, growth^42^, matrix stiffness^11^, or morphogen exposure time^28^, which are known to regulate patterning in the NT *in vivo* and NTOs *in vitro*. Our model can accommodate more complex activator-inhibitor configurations through the addition of new signaling parameters. These changes could also enable feedback from different cell types and even from neighboring organoids, as well as incorporate signal sequestration, which has been shown to play a role in ventral domain organization^42^. Given its simplicity, this model nevertheless recapitulates with remarkable fidelity the observed FP patterning phenomena. These findings underscore how the integration of *in silico* modelling, *in vitro* experimentation and transcriptomic analysis can be used as a powerful and widely applicable approach to identify the mechanisms and molecular players governing epithelial patterning.

## Acknowledgements

This work was supported by the FWO grant G087018N and FWO postdoctoral fellowship 1217220N, Interreg Biomat-on-Chip grant and Vlaams-Brabant and Flemish Government co-financing, KU Leuven grants C14/17/111 and C32/17/027 and King Baudouin Foundation grant J1810950-207421.

## Author Contributions

AA conducted experiments and analysis. AA and SG conducted FRAP experiments. AA, SG, IS and AR interpreted the data and edited the manuscript. AA and AR wrote the manuscript.

## Supplementary figures

**Supplementary figure 1.**
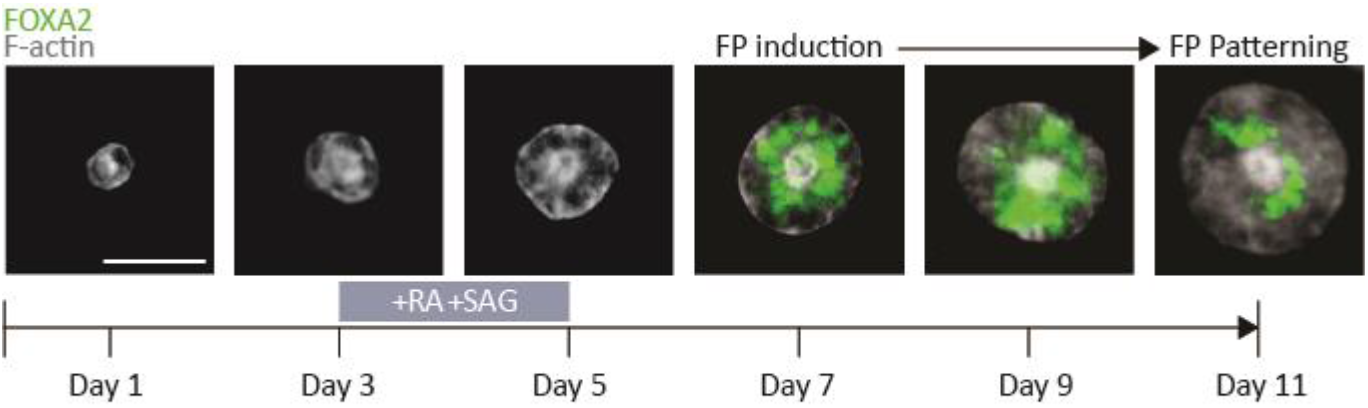
FP expression dynamics in hNTOs. Representative images of FP dynamics in hNTOs. FP induction occurs at day 7 following RA-SAG pulse. Some FP expression gradually pattern over time. Scalebar 50 μm.

**Supplementary figure 2.**
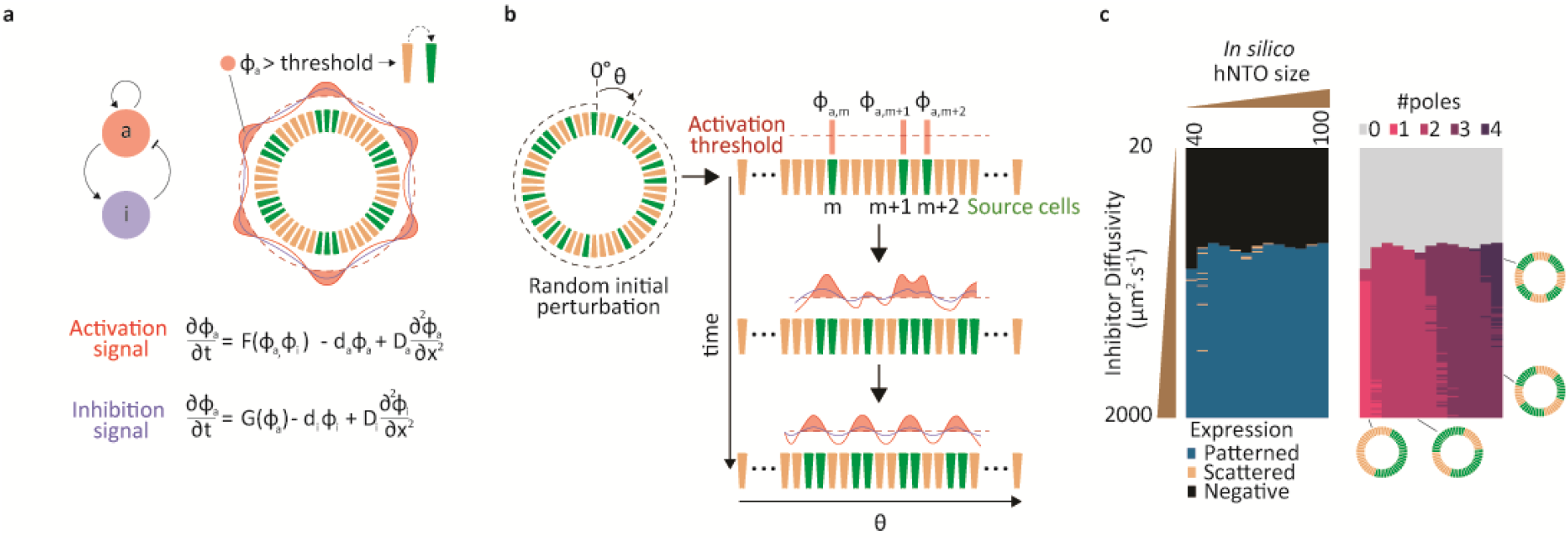
Reaction-Diffusion model for FP patterning in NTOs. **a** A 1-D two-node RD model where cells are labeled SCs (green) at periodic peaks of activation signals, Φ_a_ > threshold, where the threshold is equal to the average of the activation signal, Φ_a,avg_. **b** Initial perturbations of Φa create an instability that starts the reaction between the morphogen species eventually forcing the domain space to pattern. **c** Expression output of SCs for various inhibitor diffusion coefficients and domain sizes were the activator diffusion coefficient is 24 μm^2^/s. Associated number of poles for the same parameter configurations.

**Supplementary figure 3.**
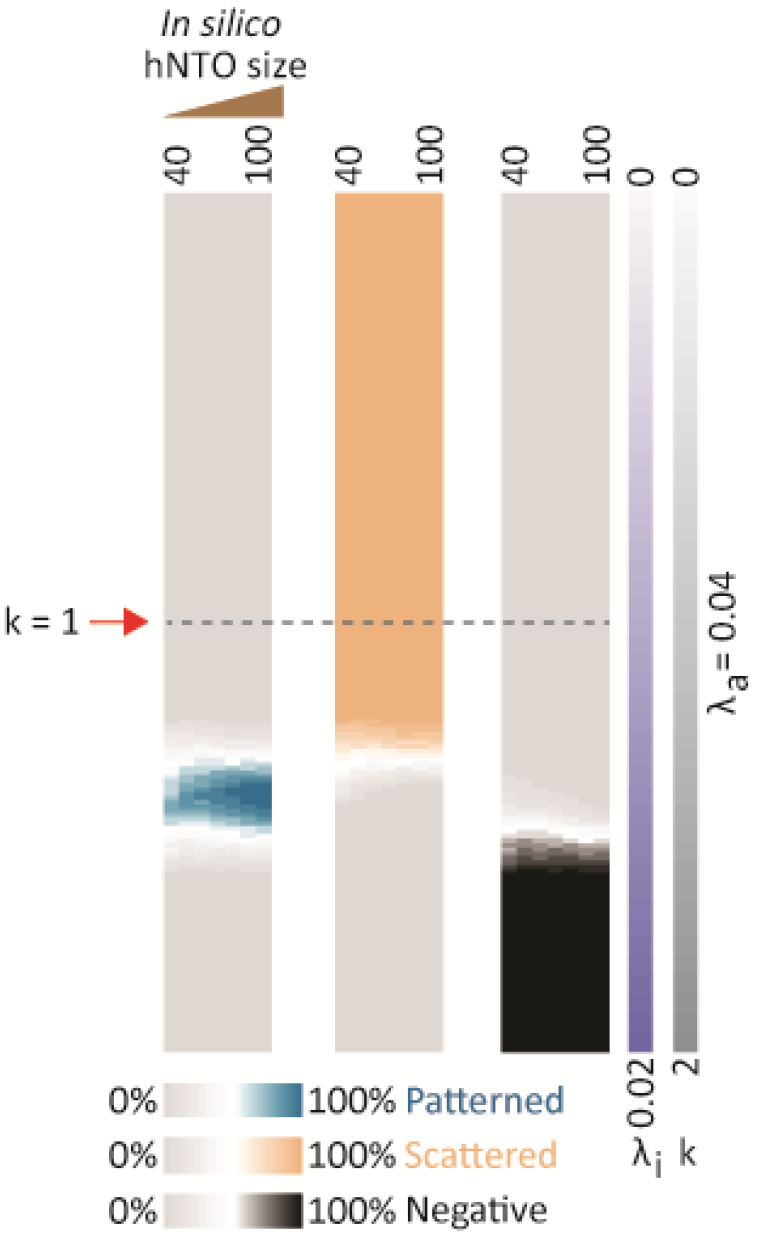
Patterned, scattered and negative *in silico* events as a function of activator and inhibitor decay constant ratio k = λ_a_/λ_i_ and domain size, where λa = 0.04. (for every simulated data point, n = 3 for a total of 1,500 *in silico* NTOs)

**Supplementary figure 4.**
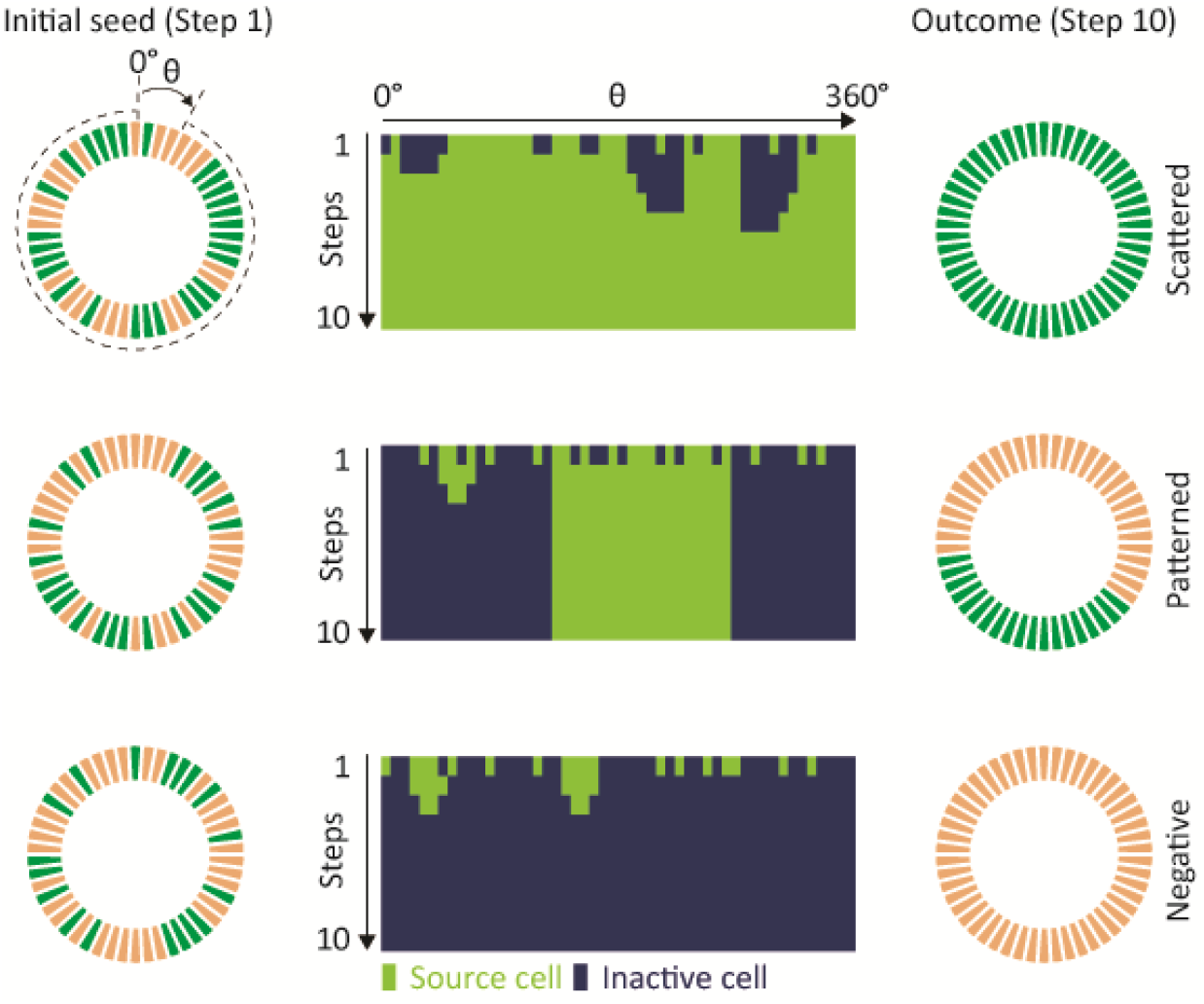
The CA model allows for expression heterogeneity. Different initial seed positions of SCs allow for scattering, patterning or negative expressions of SCs using the same parameters (k = 1.289 and λa = 0.04) and within the same domain space (46 cells).

**Supplementary figure 5.**
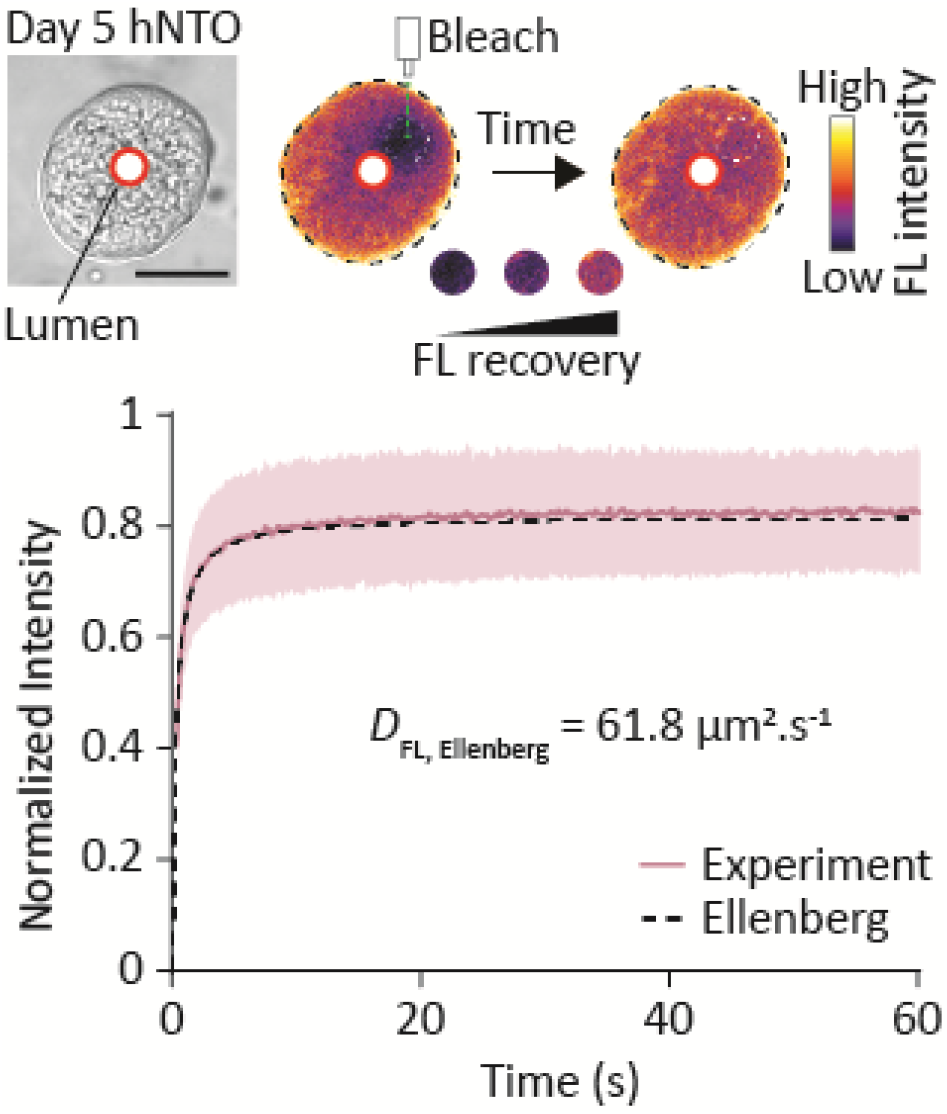
Diffusion characteristics of hNTOs matches *in vivo* conditions. A representative micrograph displaying a day 5 hNTO where the lumen region is marked with a red circle. The FL channel highlights the bleaching spot with a white dashed circle. Normalized FL intensity (solid dark purple line) and Ellenberg diffusion equation data fitting (dashed black line) as a function of time (n = 10 organoids, light purple represents SD). Scalebar 25 μm.

**Supplementary figure 6.**
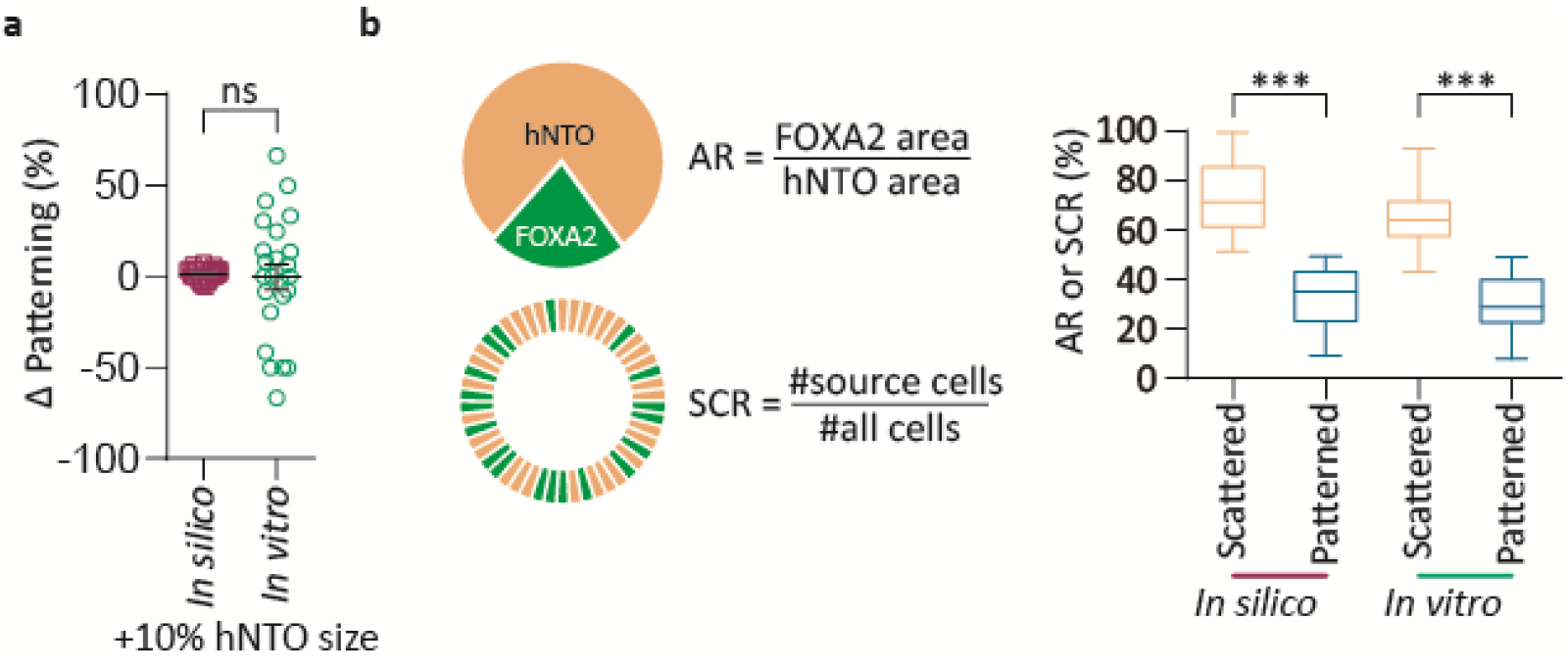
CA model predicts size-independent patterning. **a** *In silico* and *in vitro* changes in patterning for 10% increments in hNTO size or *in silico* domain size (*in silico* data from **Fig 2d**, *in vitro* data from **Fig 1d**). **b** Trends of *in vitro* ARs of day 11 hNTOs and *in silico* SCRs of final steps (n = 1,500 *in silico* NTOs and *in vitro* data from **Fig 1d**, whiskers are max and min values, hinges 25^th^ to 75^th^ percentile and horizontal line indicate the median, *** p < 0.001).

**Supplementary figure 7.**
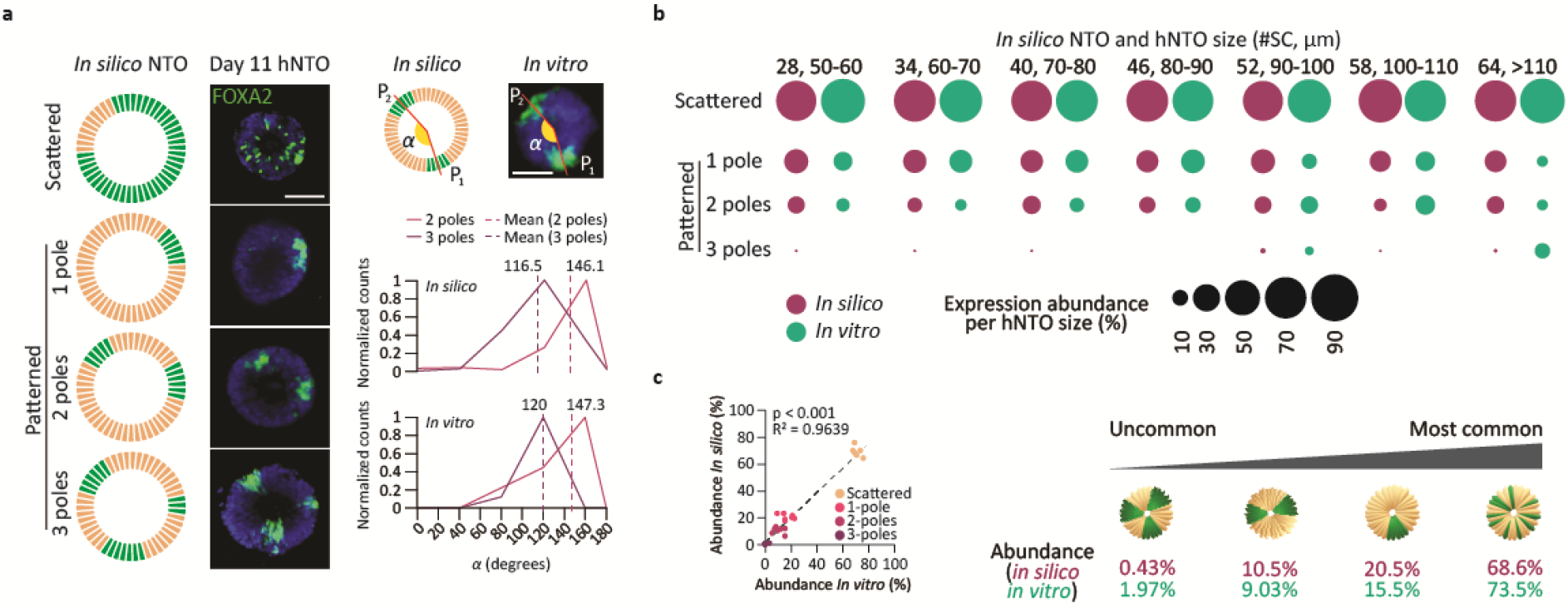
CA model accurately predicts patterning subtypes. **a** Patterning subtypes observed in *in silico* NTOs and corresponding *in vitro* observations. Normalized count of inter-pole angle α for *in silico* and *in vitro* cases (*in silico* NTOs 2-poles = 3,857, 3-poles = 219, *in vitro* hNTOs 2-poles = 15, 3-poles = 9). **b** *In silico* and *in vitro* abundance of different expression subtypes as a function of size (n = 3 for a total of 1500 *in silico* NTOs per size group, *in vitro* data from **Fig 1d**). **c** *In silico* pattern subtype correlation with *in vitro* observations (R^2^ = 0.9639). Scalebars 50 μm.

**Supplementary figure 8.**
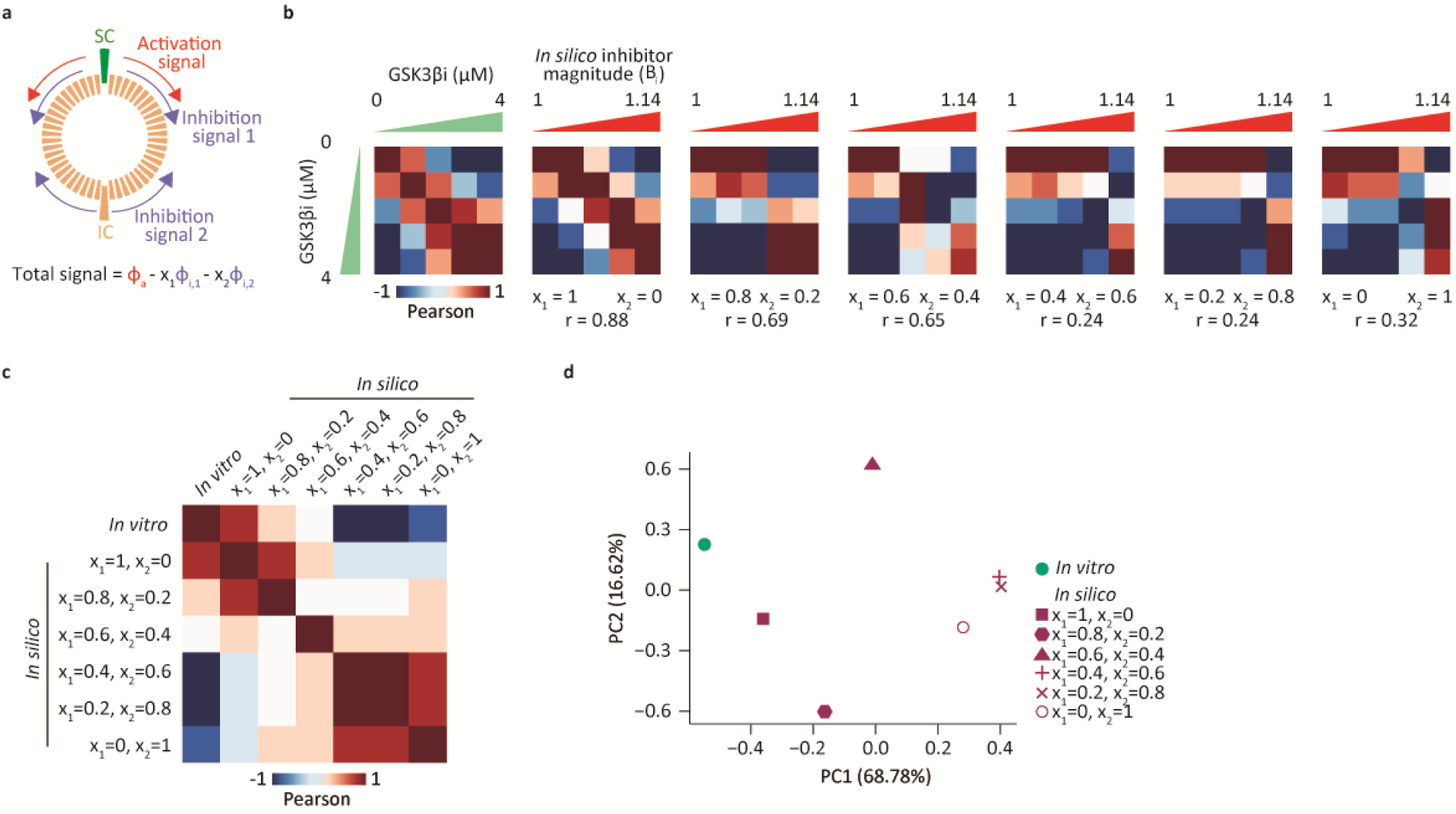
Secondary sources of inhibition do not correlate with *in vitro* data. **a** ICs participate in *in silico* signaling as secondary sources of inhibition. **b** *In silico* correlation with *in vitro* data at different SC and IC inhibition signal contributions (n = 3 for a total of 1,500 *in silico* NTOs for each *in silico* case. *In vitro* data from **Fig 4a**). **c** Summary of correlation analysis of the cases in **b**. **d** Principal component analysis of the cases in **b**.

**Supplementary figure 9.**
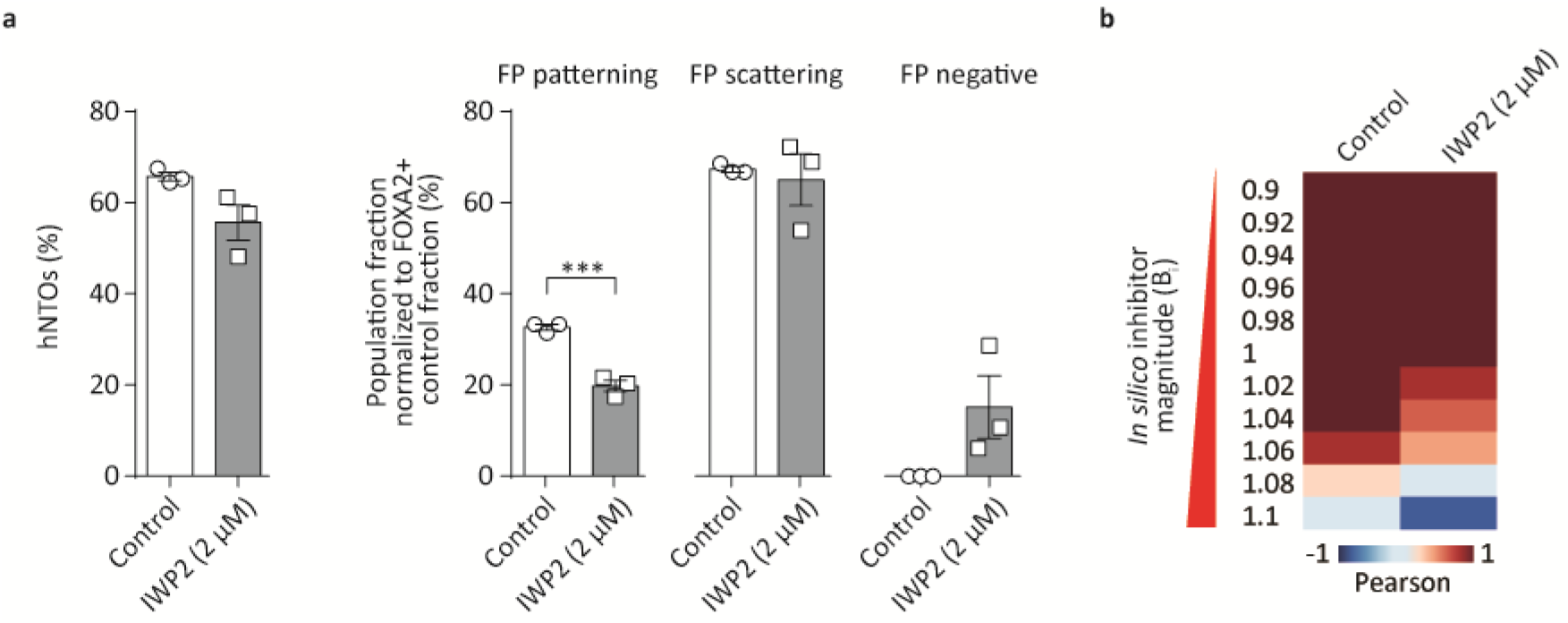
WNT inhibition reduces patterning as predicted by the model. **a** Inhibition of WNT activity by small molecule IWP2 modulates FP expression *in vitro* (n = 3 for a total of 294 hNTOs). **b** Correlation of control and IWP2 treated hNTOs with *in silico* predictions when varying inhibitor magnitude Bi (n = 3 for a total of 1,500 *in silico* NTOs per data point and *in vitro* data from a, Error bars are SD, *** p < 0.001)

## Materials and Methods

### *In silico* reaction-diffusion (RD) hNTO patterning model

We used an *in silico* 1-D RD model with one activator and one inhibitor species. The partial differential equations (PDEs) where formulated following previously defined forms^23,25^.

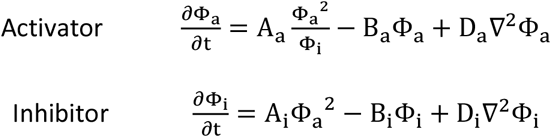

Where Φ_a_ and Φ_i_ denote the concentrations of the activator and inhibitor species respectively. The second order generation terms are defined for the activator and inhibitor as A_a_(Φ_a_^2^/Φ_i_) and A_i_Φ_a_^2^ respectively. The first order decay terms of the activator and inhibitor are B_a_Φ_a_ and B_i_Φ_i_. Values for A_a_ = 0.25, A_i_ = 1, B_a_ = 0.1 and B_i_ = 0.5, are not based on physical parameter values but are chosen to be in the range of previously used values^23^. The diffusivities of the activator and inhibitors are D_a_ and D_i_ respectively. The diffusivity of the activator molecule was set to that of SHH, a likely activator morphogen^28^ (D_a_ = 24 μm^2^/s^51,52^).

To simulate the RD model we used MATLAB (MATLAB, R2018b, The MathWorks Inc.) where a domain space was divided into 6 μm-wide units, reflecting the width of one cell. The timestep was chosen to be 1 s. For each timestep, the PDEs were solved in a forward implicit manner. The simulation is started by an initial random perturbation to the activator concentration at t = 0 s. The simulation is run for 10000 s, an adequate time for a stable pattern to form which is visually verified as non-changing over time. To evaluate patterning, cellular regions are binarized such that regions where Φ_a_ > Φ_a,avg_ are evaluated as source cells (SCs) and given a value of 1 and elsewhere as inactive cells (ICs) and given a value of 0.

To assign expression type, we adapted the scheme we used for our *in vitro* assessment. We first consider a SC domain as occupying a continuous region of SCs along the series, where ICs occupy the space (gaps) between these regions. We then evaluate the SC ratio (SCR = n(SCs) / n(total-cells)). Patterning is assigned when 1) SCR < 0.4, and 2) SCR < 0.5 but with fewer than 5 gaps. This is because upon visual inspection, we observed that for 0.4 < SCR < 0.5, SC distribution is never polarized to one side to be considered as patterned when the number of domain gaps is >5, but becomes polarized for gaps <5. For all other cases scattering is assigned. We enumerated the number of poles in our model by the number of domains detected in each simulation. By varying Di and the length of the domain space, the patterning, scattering or negative expressions and the number of poles for each conditions are used to create the heatmaps (**Supplementary Fig 2c**).

### *In silico* cellular automaton (CA) hNTO patterning model

We used an *in silico* 1-D elementary cellular automaton^27^ model to simulate patterning in hNTOs (MATLAB, R2018b, The MathWorks Inc.). Each cell is represented by a discreate element in a circular series of interacting elements. Cells are allowed two possible identities, 1) SCs, and 2) ICs. Source cells act as sources for 1) activation, Φ_a_, and 2) inhibition, Φ_i_, signals. Each signal (Φ = Be^−λx^) has a constant profile, where B is the source magnitude, λ is the exponential decay constant that controls signal decay over discreate cell positions x, and where high λ values result in rapid decay compared to lower ones. Cells are able to interact by having their respective signal profiles extend over multiple cells. The model processes 10 steps, where in each step cell interactions take place, creating a new generation of cells with modified states according to 1) activation of ICs, 2) inhibition of SCs, or 3) retention of SC and IC states. Each step advancement creates a new generation as input for the following step until the end step.

To begin the process, at step 1, cells are assigned an IC or SC identity at random, with each having equal chance of becoming either identity. Next The sums of all inhibition signals is subtracted from the sum of activation signals to obtain a net activation profile that extends over the entire cell series. A reassignment of cell states is performed and depends on the net activation value at each cell position, where values higher than a threshold *th* allows 1) ICs to assume a SC identity, or 2) SCs to retain their state. By contrast, net activation values lower than *th* 1) inactivates a SC, and 2) maintains the identity of ICs. Once all cell states have been processes, a new generation of cell series is created and passed the a new step which repeats the process. In total 10 steps are executed with the final generation evaluated for patterning using the same algorithm as in the RD model. The percentage of patterned and scattered *in silico* NTOs are evaluated per run, where a run is composed of 500 *in silico* NTOs. Three runs are evaluated (total 1,500 *in silico* NTOs) for each condition to obtain an average value for scattered, patterned and negative expression frequencies.

The *in silico* NTO sizes were evaluated by first deriving a circumference from the *in vitro* size bins (**Fig 1d**) and subsequently divide by the observed widths of nuclei (~6 μm), thereby mapping *in vitro* hTNO sizes of 50-60, 60-70, 70-80, 80-90, 90-100, 100-110 and > 110 μm to *in silico* hNTO sizes of ~28, 34, 40, 46, 52, 58 and 64 cell elements respectively.

The parameter space (**Fig 2b**), was obtained by varying Ba and Bi from 1 to 3 in increments of 0.5, and λa and λi from 0 to 0.1 in increments of 0.01 while using a domain size of 46 cells (average size after *in vitro* size mapping), producing 1,600 different parameter combinations where 157 cases result in patterning. Each condition comprised of 1,500 *in silico* NTOs where their outputs are evaluated to obtain average values for patterning, scattering and negative events. A table comprising B_a_, B_i_, λ_a_, λ_i_, B_a_/B_i_, k = λ_a_/λ_i_, patterning, scattering and negative frequencies for each condition is passed to the Seurat pipeline for data normalization and scaling^53^. Graph-based clustering was performed using the FindNeighbors function using the top 5 principle components (PCs) and the FindCluster function (resolution of 0.5) was used to group the cells with similar transcriptional profiles together. For data visualization, the dimensionality reduction technique, Uniform Manifold Approximation and Projection (UMAP), was performed with the RunUMAP function using the top 5 PCs.

Heatmaps (**Fig 2d-f**) were generated using heatmap.2 function in R. In **Fig 2d**, the domain size varied from 28 to 64 cells as per the above-mentioned *in vitro* size mapping. The activator decay constant λa = 0.04 was held constant while k was varied from 1.2 to 1.4 by varying λ_i_ while B_a_ = 2 and B_a_/B_i_ = 2. In **Fig 2e**, The domain size was held constant at 46 with λ_a_ = 0.04 and k = 1.289. The activator and inhibitor magnitudes were centered around 2 and 1 respectively and varied above and below their respective values by 0.05 magnitude points for each increment. The three expression types were then presented separately. In **Fig 2f**, the domain size varied from 28 to 64 cells as per the above-mentioned *in vitro* size mapping, with λa = 0.04 and k = 1.289. Activator magnitude was held constant at B_a_ = 2, while B_a_/B_i_ varied from 1.81 to 2.5 by varying B_i_.

The position of each pole in the patterned *in silico* NTOs was evaluated as the position of its center element. Next the distance between different pole positions was evaluated and converted to an angle (**Supplementary Fig 7a**).

### Culture medium

Essential 8 (E8) - Flex Medium Kit (ThermoFisher Scientific) was supplemented with 1% Penicillin Streptomycin (GIBCO) and used as growth medium for hiPSC culture. Neurobasal medium (GIBCO) and DMEM/F12 (GIBCO) were mixed 1:1 for the neural differentiation medium, which was supplemented with 1% N2 (GIBCO), 2% B-27 (GIBCO), 1 mM sodium pyruvate MEM (GIBCO), 1 mM glutamax (GIBCO), 1 mM non-essential amino acids (GIBCO) and 2% Penicillin Streptomycin (GIBCO).

### Human iPSC culture

Human iPSCs were cultured in Matrigel coated 6 well plates to a confluency of 60-70%, before passage every 72 h. Newly passaged colonies were exposed to Y-27632 Rock inhibitor (ROCKi) (Hellobio) at a concentration of 10 μM for the first 24 h.

### Human NTO culture in nondegradable PEG hydrogels

Human iPSC derived hNTOs were cultured in nondegrerdable polyethylene glycol (PEG) hydrogels, as previously described^20^. Briefly, hiPSCs were dissociated into single cells and embedded in a PEG hydrogel premixture. 10 μM droplets of the cell-matrix mixture were added to wells of a 96-well plate and after 20 m of gelation time, neural differentiation medium was added supplemented with 10 μM of ROCKi for the first 3 days. Retinoic acid (Stemcell Technologies) at 0.25 nM and smoothened agonist (Stemcell Technologies) at 1 μM were added to the growth medium for 2 days between days 3 and 5. This was followed by regular medium changes every 2 days until end point day 11.

### WNT perturbation experiments

Day 7 hNTOs, representing earliest FOXA2 expression, were treated with GSK3β inhibitor (Tocris, CHIR99021, 4423), an activator of WNT, until endpoint day 11 with concentrations of 1, 2, 3 and 4 μM. For WNT inhibition, IWP2 (Peprotech, 6866167-1MG) was used from day 7 at a concentration of 2 μM until endpoint day 11. Controls were treated with DMSO at dilutions similar to the highest tested concentration of GSK3βi (1:2500).

### Immunohistochemistry

Paraformaldehyde (4%) (Sigma-Aldrich) is used to fix hNTOs for 2 hours and then washed by PBS three times. Human NTOs were permeabilized and blocked using .3% Triton X (PanREAC AppliChem) and 0.5% BSA solution (Sigma-Aldrich) for 30 m. FOXA2 primary antibody mouse (Santacruz, sc-374376, 1:200) or rabbit (abcam, ab108422, 1:200) was used to stain for FP expression for 24 h and PAX6 primary antibody (DSHB, PAX6, 1:200) for more dorsal identities. This was followed by 24 h of PBS washes. For an additional 24 h, secondary antibody donkey-anti mouse Alexa Fluor 555/647 (Invitrogen) and donkey-anti rabbit Alexa Fluor 647 were used. Alexa Fluor 647 conjugated phalloidin (Abcam, 1:500) was used for filamentous actin visualization. Hoechst (1:2000) was used to nuclei visualization. Finally, this was followed by another 24 h of PBS washes.

### Imaging and image analysis

Images obtained for quantification were obtained using Zeiss Axio Observer Z1 (Carl Zeiss MicroImaging) with a Colibri LED light sources and a 10x air objective. Representative images where obtained using a Leica SP8 DIVE (Leica Microsystems) using confocal or multiphoton modes and a 25x water objective.

Pattern quantification are described elsewhere^20^. Distinguishing between 1, 2 and 3 poles relied on visual evaluation of each hNTO.

### Fluorescein recovery experiment

To *in situ* observe fluorescein diffusion within hNTOs, we first cultured hNTOs until day 5 following the hNTO protocol (**Fig 1a**). Day 5 measurements ensured capturing the ECM state at the onset of FP induction and patterning. After the RA-SAG treatment (day 5) we supplied the media with fluorescein to obtain a final concentration of 20 μM. The samples were then incubated for 30 m at 37°C and 5% CO^2^ to allow fluorescein diffusion throughout the matrix and hNTOs. We employed a confocal microscope (Leica SP8 DIVE, Leica Microsystems) to perform targeted bleaching of fluorescein molecules using 1 s burst of 480 nm laser at full power. A 10x air objective was used and a 60 s observation period of fluorescein recovery immediately followed. The bleach spot was chosen to be off-centered to avoid the acellular lumen, ensuring diffusion observations in cell-filled regions within the hNTOs. The resultant time series were analyzed using the Time Series Analyzer V3 plugin on ImageJ. For each condition, the intensity values were normalized with reference to the maximum value before bleaching. The normalized values were then averaged for each condition, and the resultant values plotted. The Ellenberg diffusion equation^29^ was used to estimate the diffusion coefficient of fluorescein.

### Single Cell RNA sequencing data processing

Data manipulation and subsequent steps were performed using the Seurat^53^ tool for single cell genomics version 3 in R version 3.4. Single Cell data were obtained from a hNTO data set^20^ that will become available at GEO. First, we subset the data to retain previously described cells with dorsal (D), intermediate (I) and ventral (V)^20^ identities summing to 1251 cells of days 5 and 11 of the hNTO differentiation protocol. The criteria for identifying dorsal, intermediate and ventral cells are described elsewhere^20^. The data was processed to find 2,000 highly variable genes using the FindVariableFeatures function. Cell cycle regression was performed. Data autoscaling was performed and reported using principal component analysis (PCA) using the RunPCA function.

### Data Clustering

Graph-based clustering was performed using the FindNeighbors function using the top 5 principle components (PCs) and the FindCluster function (resolution of 0.5) was used to group the cells with similar transcriptional profiles together. For data visualization, the dimensionality reduction technique, Uniform Manifold Approximation and Projection (UMAP), was performed with the RunUMAP function using the top 5 PCs. Cluster annotation focused on identifying D, I and V cells on the UMAPs.

### Receptor ligand interaction analysis

To systematically interrogate receptor-ligand interactions between clusters, we took advantage of the Python implementation of *CellPhoneDB* (v2.1.1)^31^. A pooled normalized count matrix containing all cells present within the scRNA-seq dataset was used as input to the algorithm. The following parameters were used: counts-data = hgnc_symbol; iterations = 1,000; threshold = 0.10. Only significant interactions (p-value < 0.05) were considered for further analysis. The receptor ligand interaction scores were ranked while highlighting those than involved known modulators of the SHH pathway, namely FGF, NOTCH, WNT, and BMP only in cases where they respectively represent more than 5% of all interactions identified per case. The Circos R package was used for visualization to display a maximum of 5 interactions per group.

### Quantification and Statistical Analysis

We used two-way ANOVA statistical tests on grouped data, and an unpaired two-tailed t-test where appropriate with a 95% confidence interval and appropriate corrections (GraphPad Prism 6, Version 6.01, GraphPad Software, Inc.). When determining patterning and scattering significance, the patterned values of the various conditions were used in statistical analysis. Similarly FOXA2+ hNTO values of the various conditions were used when determining the FOXA2+/- hNTOs statistical significance. Pearson correlations were performed to evaluate linear regression where appropriate. Statistical significance was considered for all comparisons with p < 0.05. The cor R function was used to generation the correlation maps. For hierarchical clustering and heatmap generation, we employed the R package heatmap.2.

### Data availability

This study did not generate new raw scRNAseq sequencing data. The original processed and metadata files will be made available at GEO.

